## Supplementary material for "MicroRNA-27a-5p downregulates the expression of proinflammatory cytokines in lipopolysaccharide-stimulated human dental pulp cells via NF-κB signaling pathway": PRILE checklist

### PRILE 2021

#### Checklist of items to be included when reporting laboratory studies in Endodontology*

| Section/ Topic | Item Number | Checklist Items | Reported on page number |
|---|---|---|---|
| **Title** | 1a | The Title must identify the study as being laboratory-based, e.g. “laboratory investigation” or “*in vitro*,” or “*ex vivo*” or another appropriate term | 1 |
| | 1b | The area/field of interest must be provided (briefly) in the Title | 1 |
| **Keywords** | 2a | At least two keywords related to the subject and content of the investigation must be provided | 1 |
| **Abstract** | 3a | The rationale/justification of what the investigation contributes to the literature and/or addresses a gap in knowledge must be provided | 2 |
| | 3b | The aim/objectives of the investigation must be provided | 2 |
| | 3c | The body of the Abstract must describe the materials and methods used in the investigation and include information on data management and statistical analysis | 2 |
| | 3d | The body of the Abstract must describe the most significant scientific results for all experimental and control groups | 2~3 |
| | 3e | The main conclusion(s) of the study must be provided | 3 |
| **Introduction** | 4a | A background summary of the scientific investigation with relevant information must be provided | 4~5 |
| | 4b | The aim(s), purpose(s) or hypothesis(es) of an investigation must be provided ensuring they align with the methods and results | 5 |
| **Materials and Methods** | 5a | A clear ethics statement and the ethical approval granted by an ethics board, such as an Institutional Review Board or Institutional Animal Care and Use Committee, must be described | 6 |
| | 5b | When harvesting cells and tissues for research, all the legal, ethical, and welfare rights of human subjects and animal donors must be respected and applicable procedures described | 6~7 |
| | 5c | The use of reference samples must be included, as well as negative and positive control samples, and the adequacy of the sample size justified | 6~10 |
| | 5d | Sufficient information about the methods/materials/supplies/samples/specimens/instruments used in the study must be provided to enable it to be replicated | 6~10 |
| | 5e | The use of categories must be defined, reliable and be described in detail | 6~10 |
| | 5f | The numbers of replicated identical samples must be described within each test group. The number of times each test was repeated must be described | 10 |
| | 5g | The details of all the sterilization, disinfection, and handling conditions must be provided, if relevant | 6~10 |
| | 5h | The process of randomization and allocation concealment, including who generated the random allocation sequence, who decided on which specimens to be included and who assigned specimens to the intervention must be provided(if applicable) | No |
| | 5i | The process of blinding the operator who is conducting the experiment (if applicable) and the examiners when assessing the results must be provided | No |

| | | | |
|---|---|---|---|
| | 5j | Information on data management and analysis including the statistical tests and software used must be provided | 10 |
| **Results** | 6a | The estimated effect size and its precision for all the objective (primary and secondary) for each group including controls must be provided | 10~12 |
| | 6b | Information on the loss of samples during experimentation and the reasons must be provided, if relevant | No |
| | 6c | All the statistical results, including all comparisons between groups must be provided | 10~12 |
| **Discussion** | 7a | The relevant literature and status of the hypothesis must be described | 12~14 |
| | 7b | The true significance of the investigation must be described | 13~15 |
| | 7c | The strength(s) of the study must be described | 13~15 |
| | 7d | The limitations of the study must be described | 14~15 |
| | 7e | The implications for future research must be described | 14~15 |
| **Conclusion(s)** | 8a | The rationale for the conclusion(s) must be provided | 15 |
| | 8b | Explicit conclusion(s) must be provided, i.e. the main "take-away" lessons | 15 |
| **Funding and support** | 9a | Sources of funding and other support (such as supply of drugs, equipment) as well as the role of funders must be acknowledged and described | Title Page |
| **Conflicts of interest** | 10a | An explicit statement on conflicts of interest must be provided | 1 |
| **Quality of images** | 11a | Details of the relevant equipment, software and settings used to acquire the image(s) must be described in the text or legend | 6~10 |
| | 11b | If an image(s) is included in the manuscript, the reason why the image(s) was acquired and why it is included must be provided in the text | 19~30 |
| | 11c | The circumstances (conditions) under which the image(s) were viewed and evaluated must be provided in the text | 19~29 |
| | 11d | The resolution and any magnification of the image(s) or any modifications/ enhancements (e.g. brightness, image smoothing, staining etc.) that were carried out must be described in the text or legend | No |
| | 11e | An interpretation of the findings (meaning and implications) from the image (s) must be provided in the text | 19~29 |
| | 11f | The legend associated with each image must describe clearly what the subject is and what specific feature(s) it illustrates | 19~29 |
| | 11g | Markers/labels must be used to identify the key information in the image(s) and defined in the legend | 20~30 |
| | 11h | If relevant, the legend of each image must include an explanation whether it is pre-experiment, intra-experiment or post-experiment and, if relevant, how images over time were standardised | No |

***From: Nagendrababu V, Murray PE, Ordinola-Zapata R, Peters OA, Rôças IN, Siqueira JF Jr, Priya E, Jayaraman J, Pulikkotil SJ, Camilleri J, Boutsioukis C, Rossi-Fedele G, Dummer PMH (2021) PRILE 2021 guidelines for reporting laboratory studies in Endodontology: a consensus-based development.** ***International Endodontic Journal*** **May 3. doi: 10.1111/iej.13542.**
**https://onlinelibrary.wiley.com/doi/abs/10.1111/iej.13542**

**For further details visit: http://pride-endodonticguidelines.org/prile**
